# Protocol for analysis of glycoproteomics LC-MS data using GlycReSoft

**DOI:** 10.1101/2020.05.22.108928

**Authors:** Joshua A. Klein, Joseph Zaia

## Abstract

The GlycReSoft software tool allows users to process glycoproteomics LC-MS data sets. The tool accepts proteomics database search results or a user-defined list of proteins in the sample. GlycReSoft processes LC-MS data to yield deconvoluted exact mass values. The user has the option to import a list of theoretical glycans from an external database, a curated glycan list, or a measured glycome. The tool assembles a list of theoretical glycopeptides from the lists of theoretical glycans and proteins, respectively. The program then scores the tandem mass spectra in the LC-MS data files and provides graphical views of the identified glycopeptides for each protein in the sample, and the set of glycoforms identified for each peptide sequence.

## 1. Introduction

Conventional bottom-up proteomics workflows apply most directly to identification of unmodified peptides or those with relatively small chemical or enzymatic post-translational modifications. For communication of database searching results, the mzIdentML format was developed (1,2) and includes such small PTMs that fit with the original proteomics use cases. Complex glycosylation does not fit within the scope of bottom-up proteomics as originally defined. One problem is that glycosylation, resulting from ER and Golgi-mediated biosynthetic reactions, is heterogeneous as a rule, necessitating the definition of a range of post-translationally modified forms of a given modified peptide sequence. Another problem is that the glycans undergo dissociation during the tandem MS experiment, and there is no way to specify the product ions using existing data standards.

The proteoform concept was formulated in response to the observation of protein heterogeneity using top-down MS (3,4). Proteoforms, as different modified forms of a given gene product, diversify the functions that can be attributed to a protein through allosteric regulation and/or activation of interactions with binding partners or adapter molecules. For complex glycosylation, it is not uncommon to observe more than 30 glycoforms at a glycosite. The number of theoretical glycoforms thus multiplies as the number of protein glycosites increases, rapidly exceeding the number of proteoforms that could be made by a cell (5). Therefore, the goal of glycoproteomics is to identify the subset of the theoretical glycoforms that exist in a given biological context.

As summarized in recent reviews, glycopeptides are assigned in mass spectrometry experiments using features including mass, elemental composition, tandem mass spectra, and time of elution or migration from the separation system (liquid chromatography or capillary electrophoresis) (6-8). In principle, the confidence of an assignment improves as the mass spectrometer accuracy increases. For the tandem MS step, three types of product ions define the glycopeptide in terms of peptide sequence, glycan composition and glycosylation site(s). Glycopeptides dissociate to form low mass oxonium ions corresponding to saccharide fragments. Neutral losses of saccharide units give rise to a series of peptide + Y_n_ ions. The presence of peptide backbone product ions is particularly useful for identifying the peptide sequence. Collisional dissociation defines the peptide sequence and glycan composition. In favorable cases, the site of glycosylation can be defined for singly glycosylated peptides. For multiply glycosylated peptides, collisional dissociation defines the total glycan composition but often does not define the glycosylation at individual peptide sites. Electron activated dissociation methods produce preferential cleavage of the peptide backbone and are therefore more likely to succeed in assigning multiply glycosylated peptides.

GlycReSoft is an open-source software program for processing glycomics and glycoproteomics LC-MS data (9,10) (Figure 1, see Note 1). The program uses a deconvoluter based on DeconTools (11) and MD-DeconV (12) that employs both peptide and glycopeptide averagine values that identifies glycopeptide precursor ion elemental compositions based on isotope pattern matching. The program uses the list proteins identified using a proteomics search engine in the form of an export in mzIdentML or FASTA format. The user can specify the range of theoretical glycan compositions by importing from an external database or defining a theoretical list from algebraic combinations of glycans. The program also accepts measured glycan profiles from experimental data. Users can also import curated glycan lists. The program then calculates the theoretical glycopeptide precursor ions using the theoretical glycan lists and the observed sample proteome.

**Figure 1.**
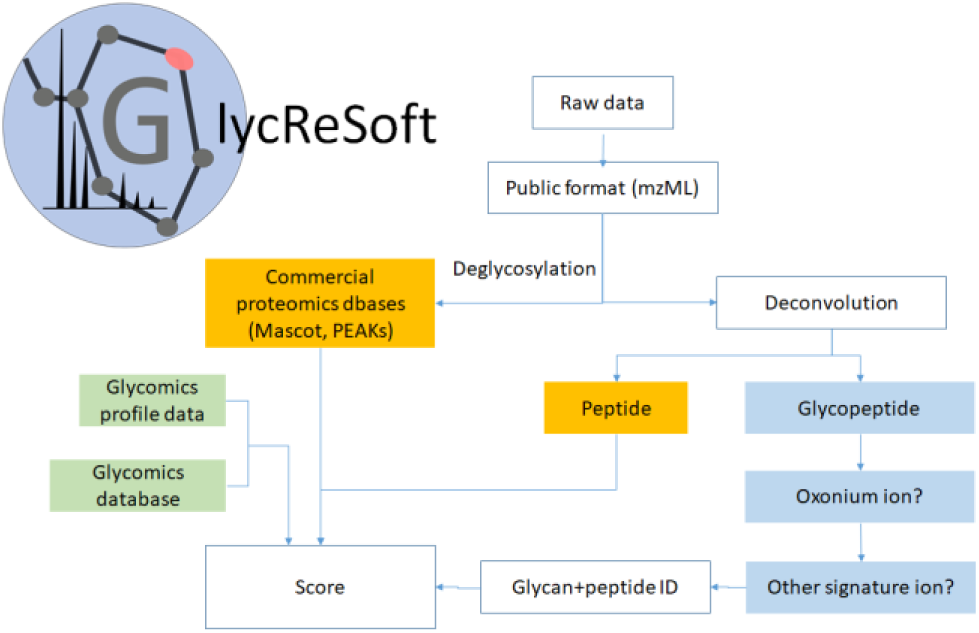
Glycresoft pipeline for processing glycopeptide LC-MS data

We provide here a method for using GlycReSoft for analysis of a human α1-acidglycoprotein tryptic digest from a research publication (13). For this, the user is directed to download data files from the Pride archive. These consist of the reversed phase LC-MS data acquired using a Thermo-Fisher Scientific Q-Exactive plus mass spectrometry system, see Figure 2.

**Figure 2.**
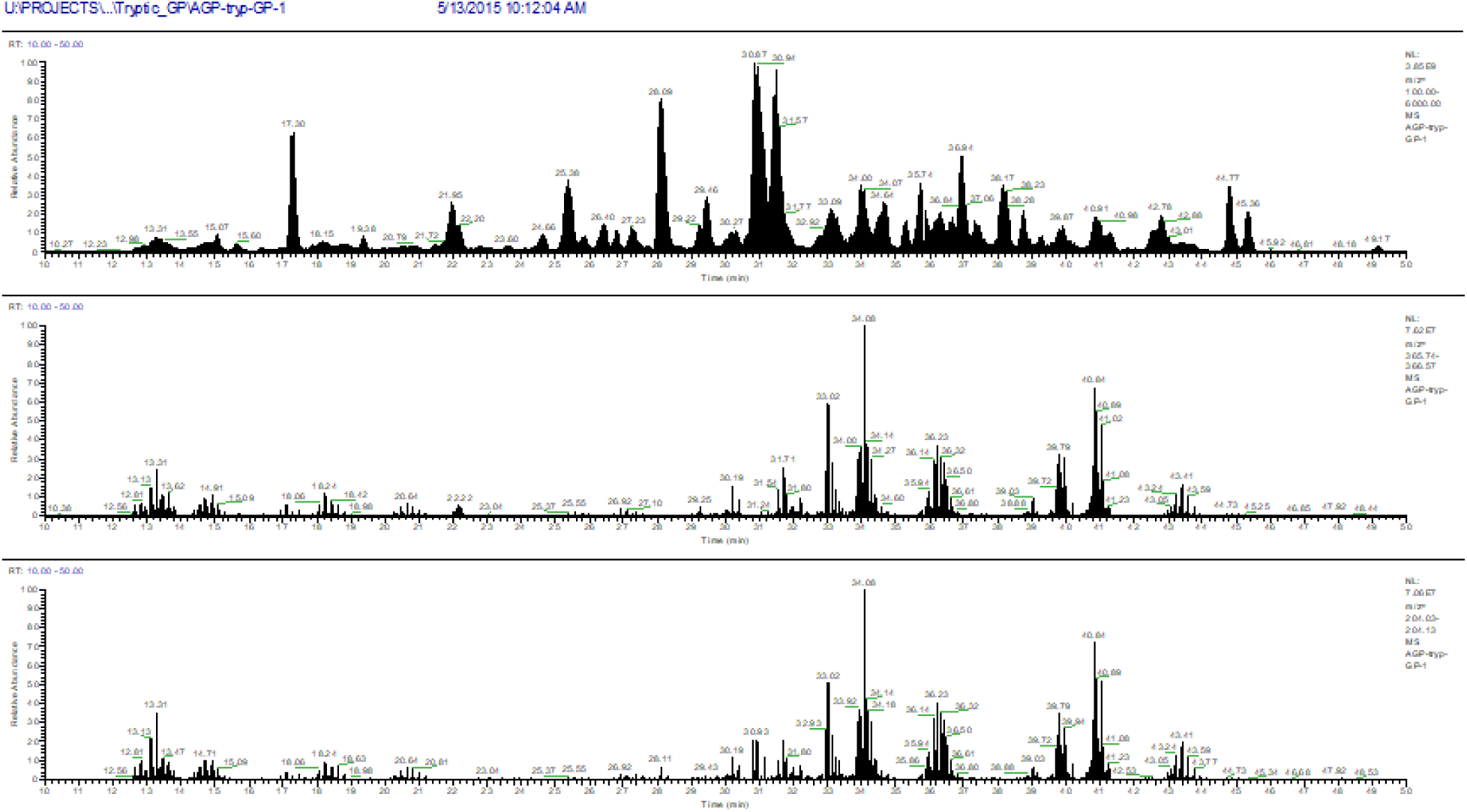
Human α1-acidglycoprotein tryptic peptides analyzed using reversed phase LC-MS (13). The top panel shows the extracted total ion chromatogram. The middle panel shows the extracted ion chromatogram (EIC) for the Hex-HexNAc oxonium ion. The bottom panel shows the EIC for the HexNAc oxonium ion.

## 2. Materials

### 2.1. Installation

2.1.1. Windows. Download and install the Windows Graphical Installer (14).

GlycReSoft will prompt the user to create a working folder in which results files will be stored.

2.1.2. LINUX. A Windows command line interface is also available along with technical documentation (14).

### 2.2. Quick tour

GlycReSoft implements algorithms for:

- Generation of lists of theoretical glycan compositions using algebraic rules, the glySpace database network (15), or from text files;
- Generation of lists of theoretical glycopeptide masses using protein sequences in FASTA format or proteomics search results in Human Proteome Organization (HUPO) Proteome Standards Initiative (PSI) (16) mzIdentML format (1,2), combined with a glycan search space;
- Support for *N*-linked, *O*-linked, or GAG-linker glycopeptides;
- Deisotoping and charge state deconvolution of glycan and glycopeptide mass spectra;
- Identification and quantification of glycans by MS and glycopeptides by MS/MS.

GlycReSoft reads LC-MS data in mzML (17) and mzXML (18) formats. It also reads Thermo-Fisher RAW file format. It is available as a Windows precompiled build (described here), compatible with Windows 8 and 10, and a UNIX/LINUX command line interface (14).

## 3. Methods

3.1. Download public proteomics and glycoproteomics data on human alpha-1-acid glycoprotein digests (13). File names indicate protease used and whether or not samples were deglycosylated using peptide N-glycosidase F (PNGaseF) before analysis. The files are available through the following PRIDE repository (19):

Project Name: Influenza A virus-integrated glycomics, proteomics and glycoproteomics

Project accession: PXD003498

Project DOI: 10.6019/PXD003498

Keys to data:

Tryp-tryptic digest

Chymo-Chymotryptic digest

O16-Samples subjected to PNGaseF deglycosylation in presence of regular water (H216O).

GP-No deglycosylation.

**3.1.1.** Download the files “AGP-tryp-GP.raw” and “AGP_O16_tryp_peptides_1_1_0_new.mzid.gz” (see Note 2).

**3.1.2.** Convert the “AGP-tryp-GP.raw” file to mzML format using ProteoWizard MS_convert (20) (see Note 4) using the default parameters (Figure 3).

**Figure 3.**
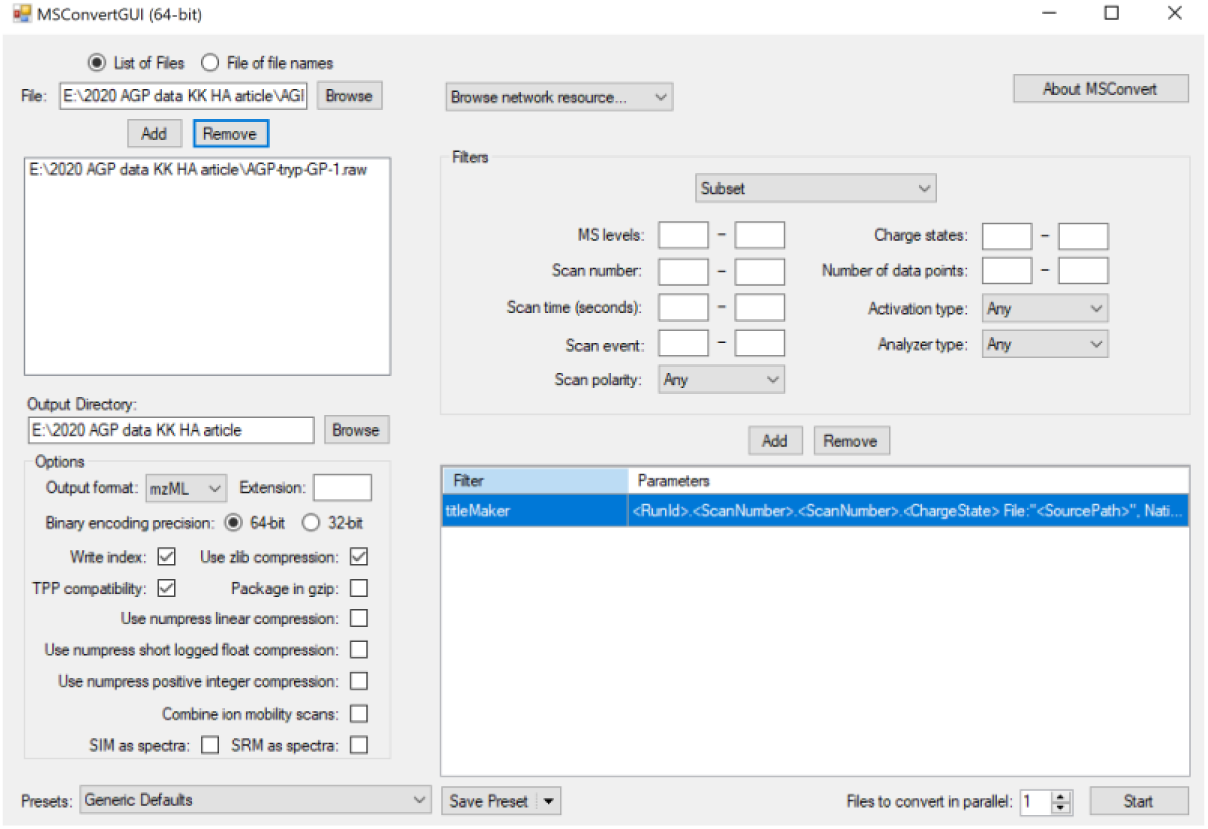
Screen shot of the MS_convert utility

### 3.2. Create a glycan search space

Click on “BUILD A GLYCAN SEARCH SPACE” (Figure 4). In the window, give the glycan search space a name by clicking on “Hypothesis Name”. Select Reduction type “native” and Derivatization Type “native”. Click “COMBINATORIAL HYPOTHESIS” and use the default monosaccharide and algebraic rules (see Note 3). Click “GENERATE”. The bar on the left indicates the task status.

**Figure 4.**
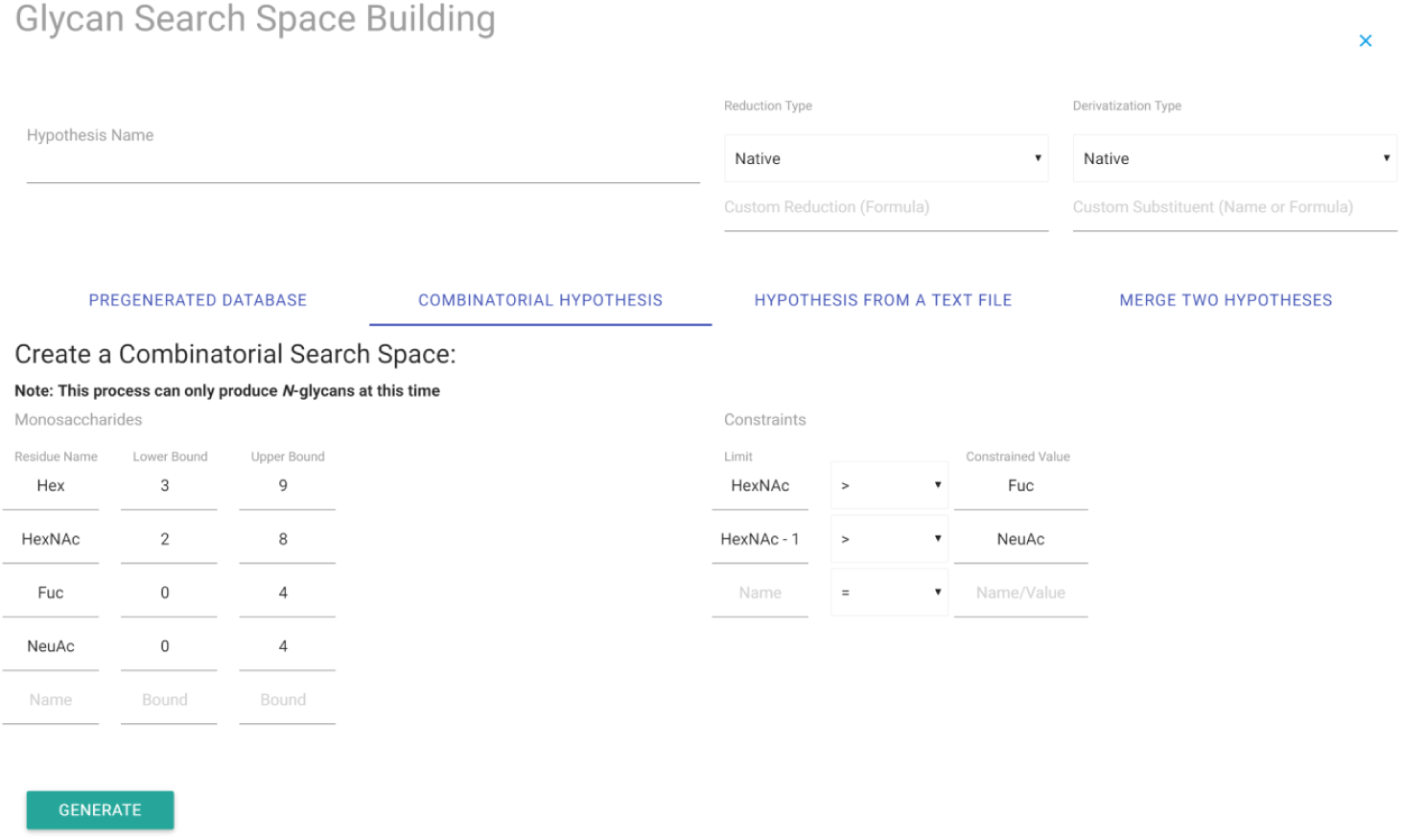
Glycan Search Space Building window

### 3.3. Create a glycoproteomics search space

Click “BUILD A GLYCOPEPTIDE SEARCH SPACE” (Figure 5). Click “Hypothesis Name” and enter a name. Click “SELECT FILE”, then click on the protein list file “AGP_O16_tryp_peptides_1_1_0_new.mzid”. Under “Select a Glycan Hypothesis or Sample Analysis” select the name of your glycan hypothesis. Leave all other parameters at their default values (see Note 5). Click “GENERATE”. The bar on the left indicates the task status.

**Figure 5.**
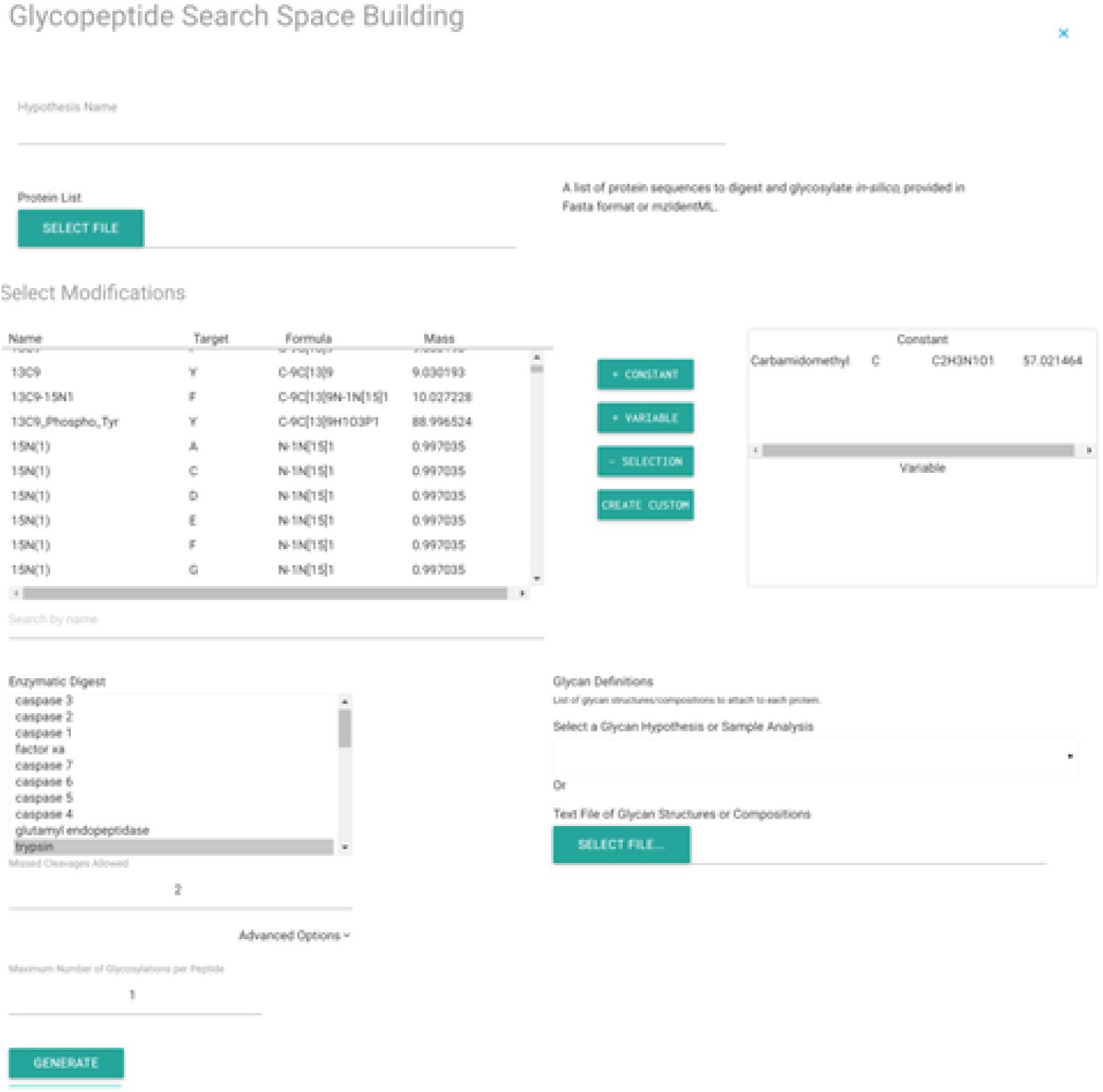
Glycopeptide Search Space Building window

### 3.4. Run LC-MS preprocessing steps

Click “ANALYZE SAMPLE”. Click “SELECT MZML FILE” (Figure 6). Provide a sample name. Under “Preset Configurations” select “LC-MS/MS Glycoproteomics”. Under “MS1 Parameters” “Averagine” select “glycopeptide”. Under “MSn Parameters” “Averagine” select “glycopeptide”. Use the default settings for all other parameters. Click “SUBMIT”. The bar on the left indicates the task status.

**Figure 6.**
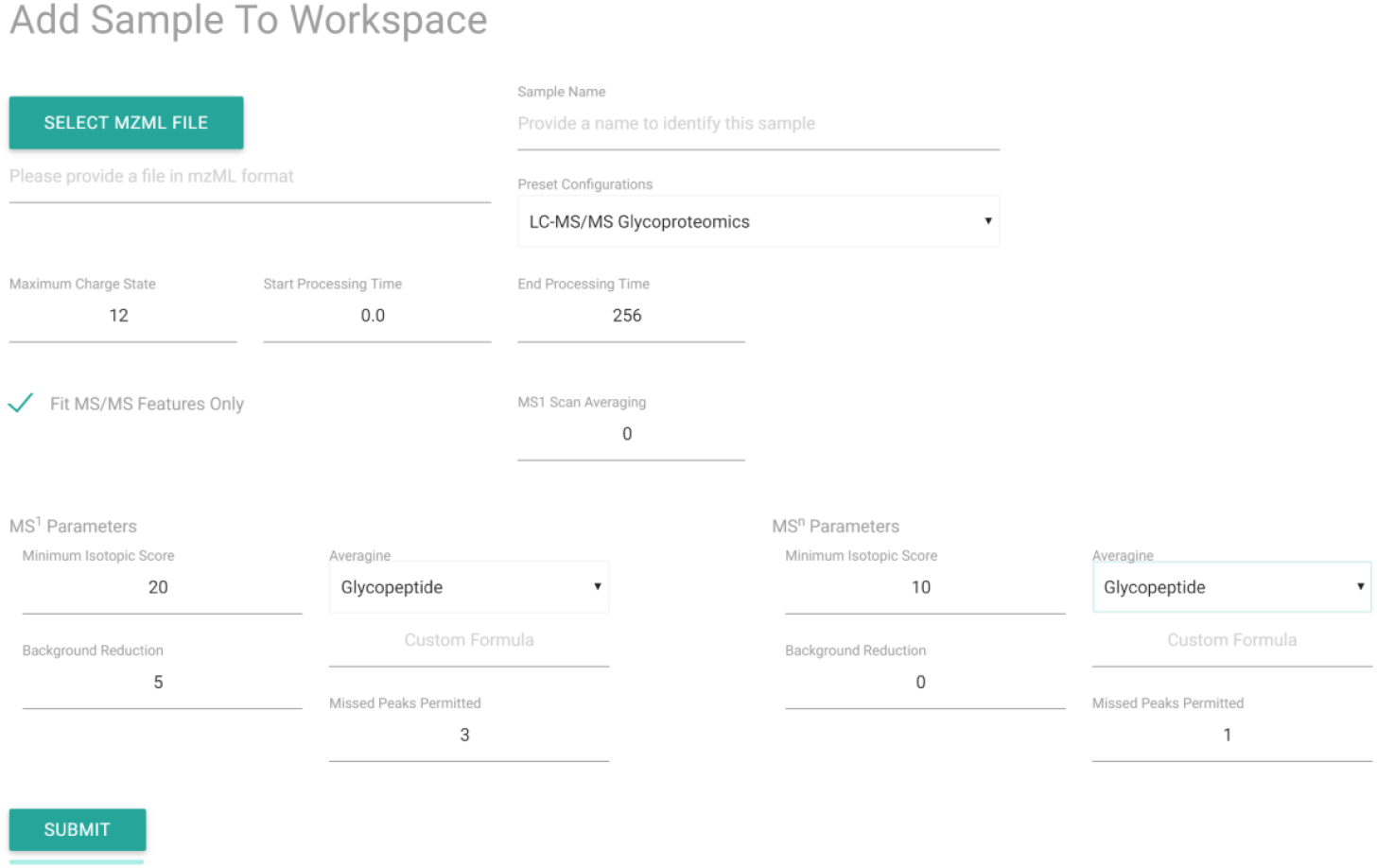
ANALYZE SAMPLE window

### 3.5. Search glycopeptide sequences

Click “SEARCH GLYCOPEPTIDE SEQUENCES” (Figure 7). Select a sample from the menu under “Select One or More Samples”. The samples correspond to pre-processed mzML data files stored in the working directory. Under “Choose a Hypothesis” select the name of the glycopeptide search space file created in step 3.3. Leave all other parameters as their default values. Click “SUBMIT”. Progress is shown on the task window on the left side of the screen.

**Figure 7.**
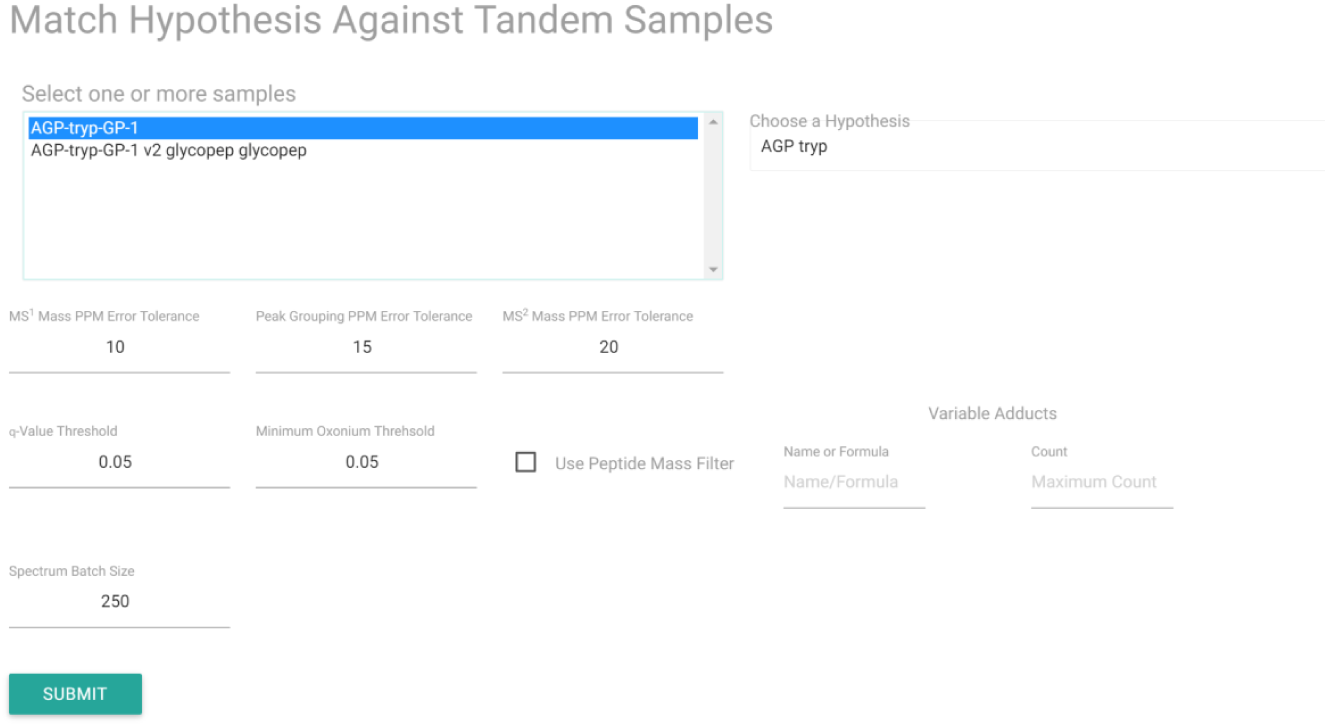
Search Glycopeptide Sequences window

### 3.6. View results

Under “Analyses” click on the results that you wish to view. Clicking on the “OVERVIEW” tab, GlycReSoft displays results for each protein in the proteome. Clicking on the “GLYCOPEPTIDES” tab displays a table of all glycopeptides detected for the selected protein (Figure 8). Clicking on individual rows jumps to display of the extracted ion chromatogram, tandem mass spectrum, and product ion table. Scrolling down, the program displays a pileup diagram of the glycopeptide glycoforms identified for each glycosite in the protein sequence. Mousing over the pileup diagram displays glycopeptide mass, sequence, glycan composition and MS2 statistics. Clicking on an individual glycopeptide bar will display the extracted ion chromatogram, tandem mass spectrum, and product ion table (Figure 9). Clicking on the “GLYCOPEPTIDES” tab displays a table of all glycopeptides identified (Figure 10). Clicking on the “SITE DISTRIBUTION” tab displays bar plots of the glycoforms identified for each peptide sequence (Figure 11).

**Figure 8.**
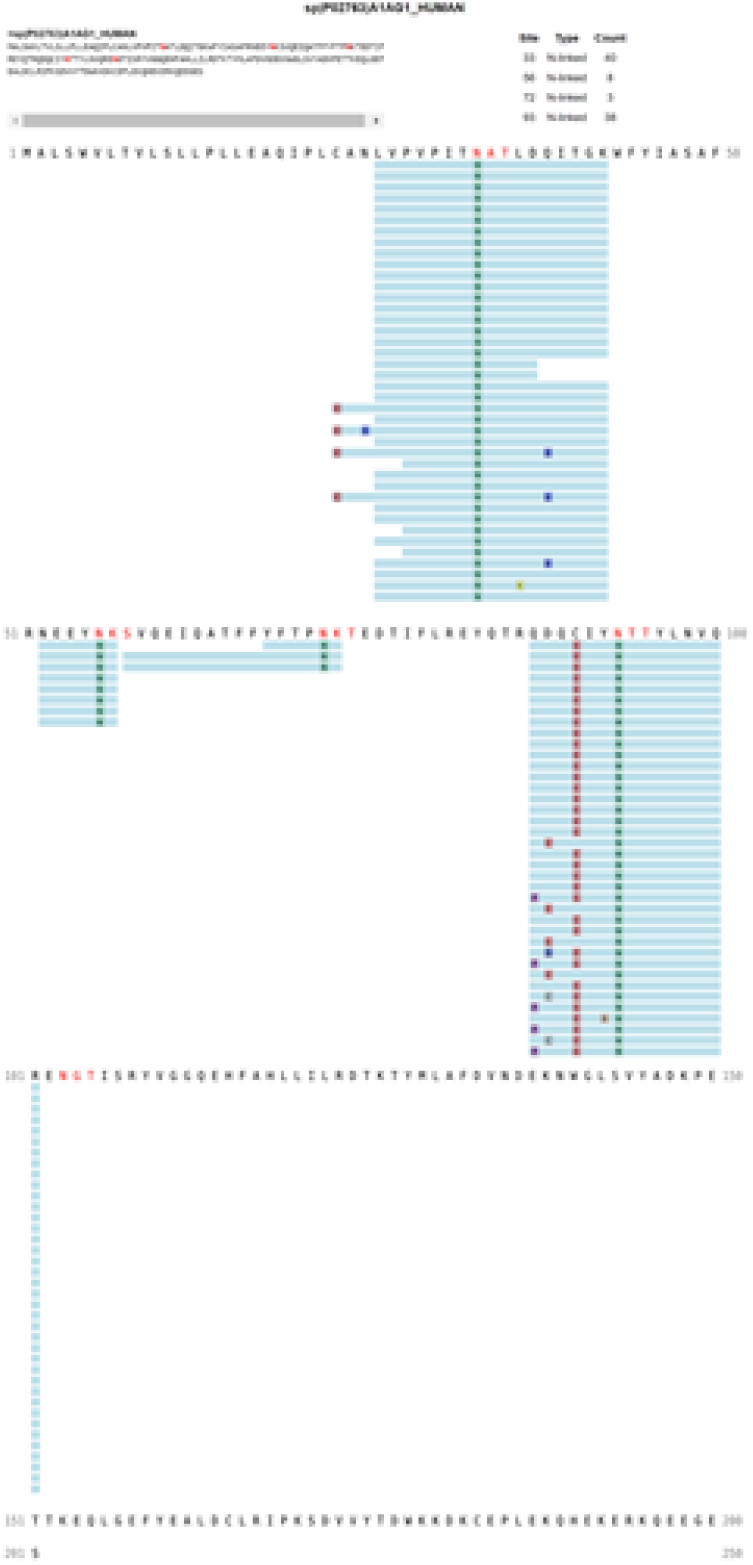
Glycopeptide pileup diagram for AGP1.

**Figure 9.**
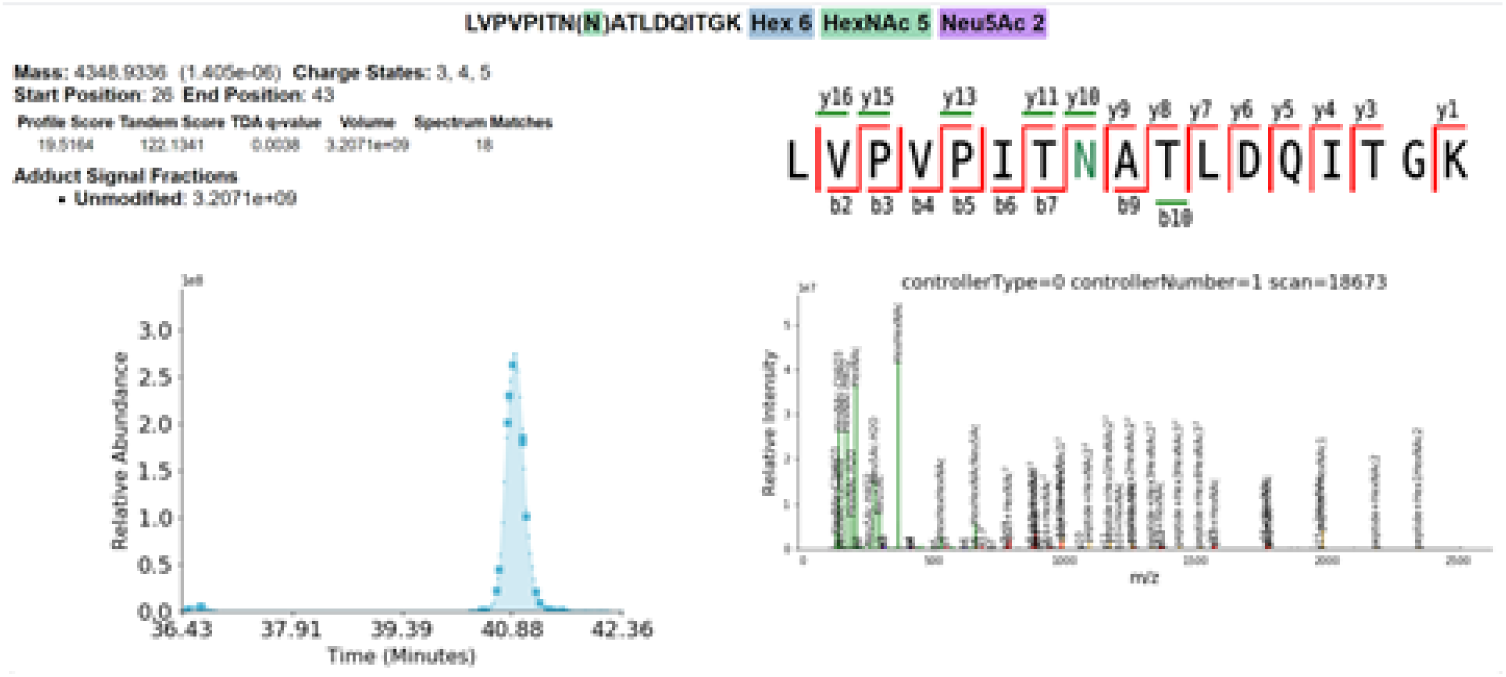
Glycopeptide extracted ion chromatogram, annotated tandem mass spectrum, fragmentation diagram, and tandem MS statistics.

**Figure 10.**
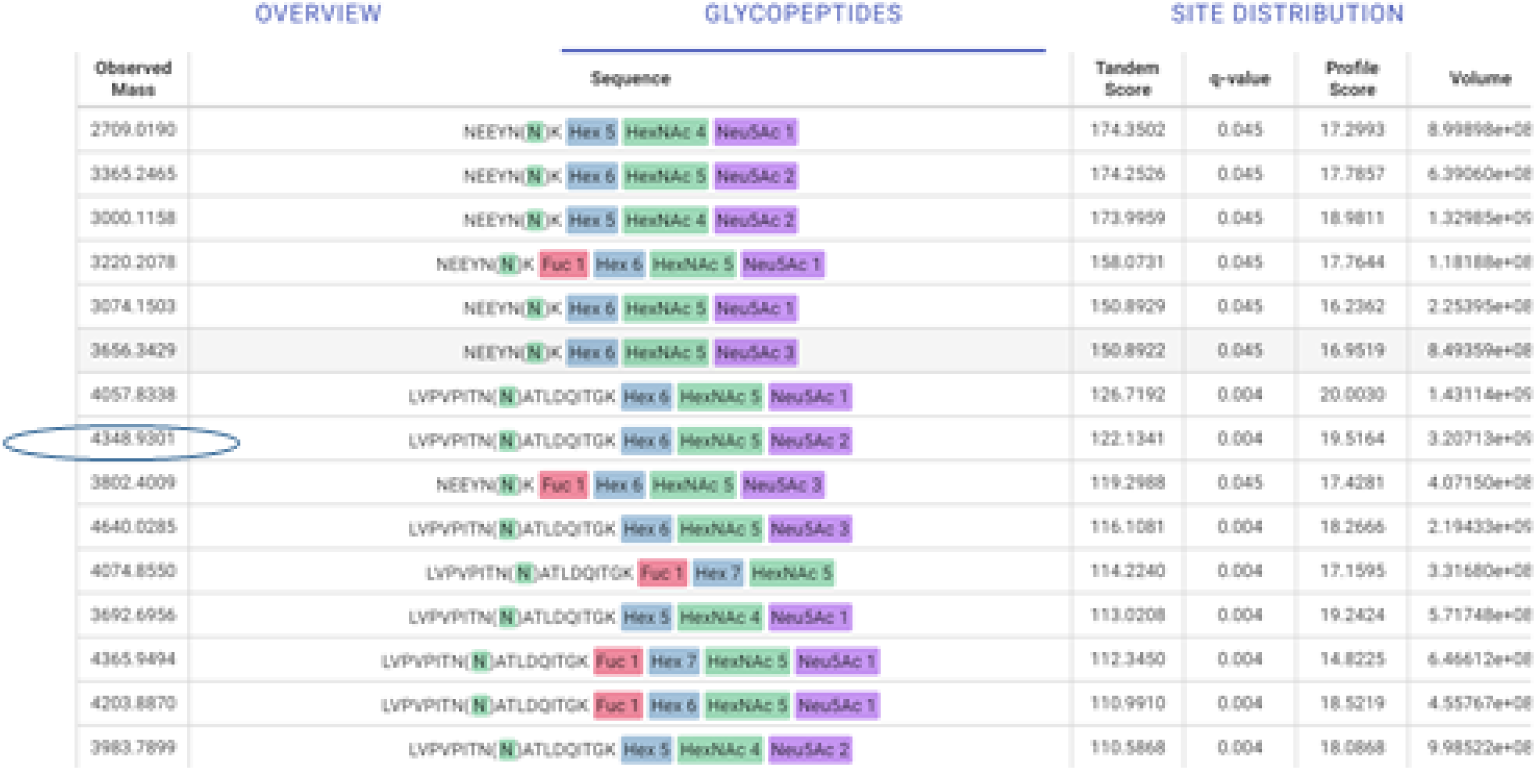
Example table of identified glycopeptides (partial listing)

**Figure 11.**
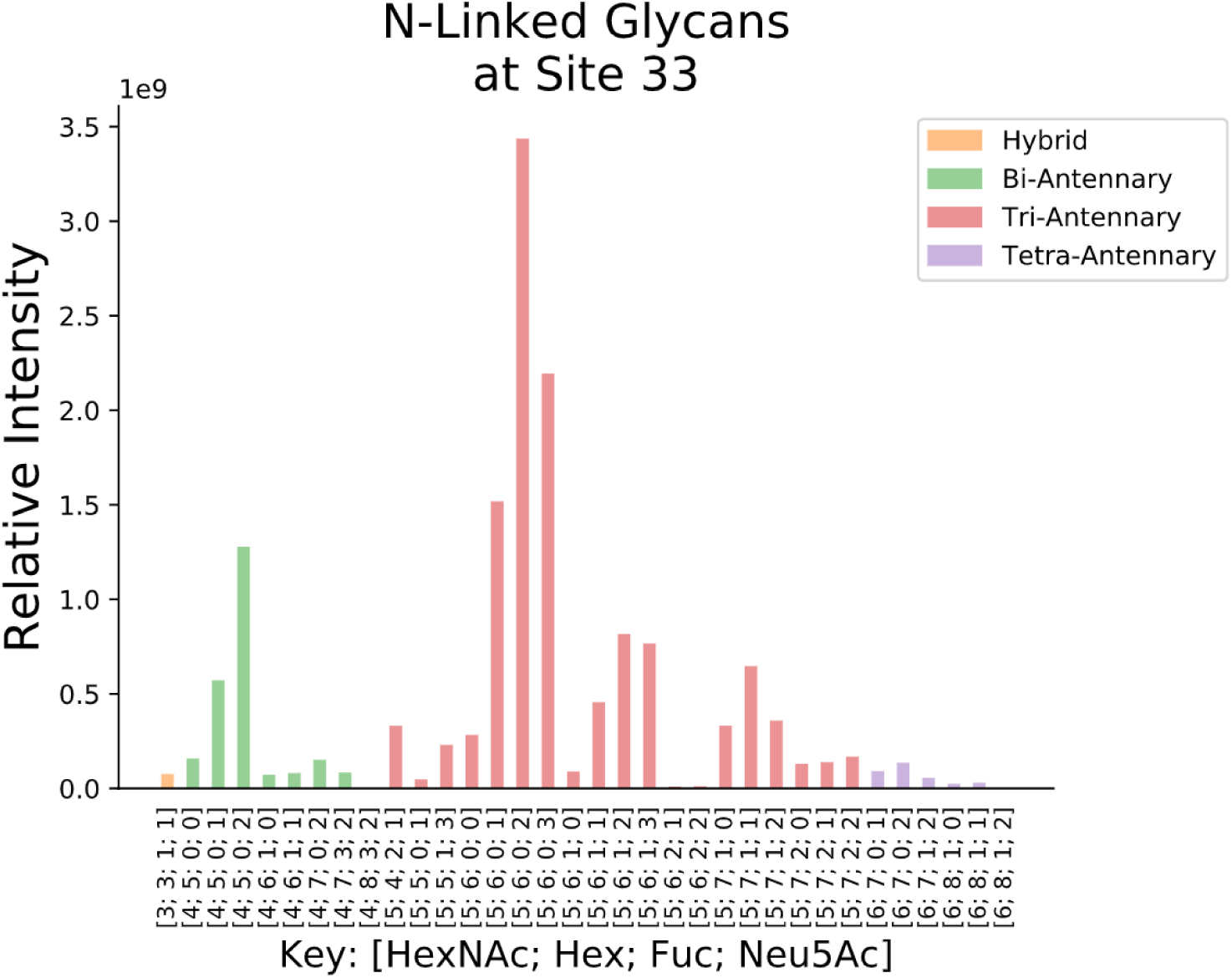
Example bar plot showing all glycoforms identified for a peptide sequence.

### 3.7. Export results

Click on the disk icon to display the results export options. GlycReSoft exports results to the as comma separated value files. It generates annotated pdf files for each tandem mass spectrum. It generates a HTML report that is viewable using a web browser.

## 4. Notes

Note 1. The use of GlycReSoft for glycoproteomics is described here.

Note 2. Unzip the *.gz directory using 7-Zip (https://wcww.7-zip.org/).

Note 3. The monosaccharide list can be modified to include NeuGc or other monosaccharides as desired.

Note 4. The ProteoWizard package is available for download at: http://www.proteowizard.org/download.html

Note 5. When using mzIdentML files the fixed modifications, proteases, and number of missed cleavages are read from the mzIdentML file.

Note 6. For MSn spectra, the peptide averagine works best higher normalized collision energy (>25) dissociation. The reason is that most b/y ions are almost entirely peptide, and the peptide+Y ions dominated by the intact peptide most of the time. When lower normalized collision energy is used, peptide+Y ions with more glycan are present, when a combination of peptide and glycopeptide averagines works better.

## 5. Acknowledgements

The development of GlycReSoft was supported by NIH grants P41GM104888 and U01CA221234.

## Notes

### Competing Interest Statement

The authors have declared no competing interest.

